# Encoding the Space of Protein-protein Binding Interfaces by Artificial Intelligence

**DOI:** 10.1101/2023.09.08.556812

**Authors:** Zhaoqian Su, Kalyani Dhusia, Yinghao Wu

## Abstract

The physical interactions between proteins are largely determined by the structural properties at their binding interfaces. It was found that the binding interfaces in distinctive protein complexes are highly similar. The structural properties underlying different binding interfaces could be further captured by artificial intelligence. In order to test this hypothesis, we broke protein-protein binding interfaces into pairs of interacting fragments. We employed a generative model to encode these interface fragment pairs in a low-dimensional latent space. After training, new conformations of interface fragment pairs were generated. We found that, by only using a small number of interface fragment pairs that were generated by artificial intelligence, we were able to guide the assembly of protein complexes into their native conformations. These results demonstrate that the conformational space of fragment pairs at protein-protein binding interfaces is highly degenerate. Our study illustrates how artificial intelligence can be used to understand and characterize protein-protein binding interfaces. The method will be potentially useful to search for the conformation of unknown protein-protein interactions. This result demonstrated that the structural space of protein-protein interactions is highly degenerate under the representation of interface fragment pairs. Features in this degenerate space can be well characterized by artificial intelligence. In summary, our machine learning method will be potentially useful to search for and predict the conformations of unknown protein-protein interactions.

## Introduction

Proteins rarely act alone in a highly crowded cellular environment [1]. Their biological functions depend closely on the partners they interact with [2]. The dynamics of protein-protein interactions are largely determined by the structural properties of their binding interfaces [3]. It has been found that mutations of residues at the interface of a protein complex can significantly change its stability and thus affect its cellular function [4]. Therefore, deciphering the structure of how proteins interact with each other is a crucial step towards understanding the functions of these molecules. The traditional techniques that can be used to experimentally determine the structure of protein complexes include x-ray crystallography, NMR spectroscopy and electron microscopy [5]. The rate of solving new structures by these experimental methods is about 200–500 protein complexes per year [6]. By examining the sequence space of protein complexes, it was estimated that the total number of unique interaction types between proteins is on the scale of 10,000 [7]. As a result, at the current rate of structure determination, it would take at least two decades before a complete set of protein complex structures is available. This reality highlights the urgent need for developing efficient computational methods that are able to accurately predict the structure of protein complexes, especially when the structures of homologous proteins are not available.

The original endeavor in structural modeling of protein-protein interactions has been focused on rigid-body docking. Docking methods explore all possible binding modes of two proteins without a priori knowledge of their complex structures [8-10]. As a result, their success often depends on the size and shape complementarity of the interface area, and the hydrophobicity of interface residues. Another challenge of docking is the limitation of sampling the entire conformational space. Different from docking, other methods use structurally characterized complexes as templates to construct models of unknown protein complexes [11-16]. However, template-based modeling is essentially limited by the range of protein complexes that can be modeled, due to the fact that the current complex structure library is far from complete in covering the quaternary structure space of nature. Computational biology has greatly benefited from recent advances in artificial intelligence (AI), especially deep learning [17, 18]. The application of these algorithms to structural biology has drawn enormous attentions [19, 20]. For instance, AlphaFold has gained tremendous success in protein tertiary structure prediction [21]. It has recently been extended to model the structure of protein complexes, which is called AlphaFold-multimer [22]. Other similar deep-learning-based methods, such as RoseTTAFold [23], have also achieved high accuracy in predicting the structures of multimeric protein complexes.

The successes of these deep-learning-based methods in protein complex modeling originate from previous observations that the binding interfaces of diverse protein-protein interactions could be highly correlated [11, 24]. It was previously found that 80% of the interfaces in protein complexes could form a densely connected network [25]. Moreover, we previously defined an interface fragment pair as a pair of 9-residue-long fragments from each side of a dimeric protein complex [26]. We collected all interface fragment pairs from a structure database of protein complexes and grouped them together based on structural similarity. We showed that, by only using a small number of highly representative interface fragment pairs, we can reconstruct the native-like quaternary structures for most of the protein complexes in a large-scale protein docking benchmark. Our results suggested that the structural space of protein-protein interactions can be decomposed by a limited number of fragment pairs at their binding interfaces.

In this work, we further used machine learning to study interface fragment pairs. Among various machine-learning-based methods, autoencoders have been used to sample the conformational space of both structured and intrinsically disordered proteins [27, 28]. We first developed a generative autoencoder to represent interface fragment pairs in a low-dimensional latent space, and reconstruct their structures back in Cartesian coordinates. Samples encoded in the latent space were further clustered and visualized by a self-organizing map (SOM) [29]. After training, we used the autoencoder to generate new conformations of interface fragment pairs through SOM-weighted sampling. We found that most newly generated interface fragment pairs are close to native-like structures. They can guide the assembly of protein complexes. These results demonstrate that the conformational space of fragment pairs at protein-protein binding interfaces is highly degenerate. Our study illustrates how artificial intelligence can be used to understand and characterize protein-protein binding interfaces. The method will be potentially useful to search for the conformation of unknown protein-protein interactions.

## Methods

### Dataset of fragment pairs at protein-protein binding interfaces

Given a pair of interacting proteins *I* and *J*, we first find all residues that form inter-molecular contacts at its binding interface. Specifically, for a residue from protein *I* and another residue from protein *J*, we calculated the distances of all side-chains atoms between these two residues. If at least one of these inter-residue atomic distances are smaller than the cutoff (5 Angstrom), these residues are thought to form an inter-molecular contact. An interface fragment pair was further defined based on the inter-molecular residue contact by taking account of the information of their local backbone conformation. Each fragment is represented by a window of 9 amino acids that is centered at the corresponding interface residue. The local conformation of the fragment is described by the coordinates of their Cα atoms. As a result, two fragments centered at an inter-molecular residue contact form an interface fragment pair, as shown in **Fig 1a** The Cartesian coordinates of all 2×9=18 Cα atoms are used to represent the structure of an interface fragment pair. Therefore, it contains 3×18=54 degrees of freedom.

**Figure 1:**
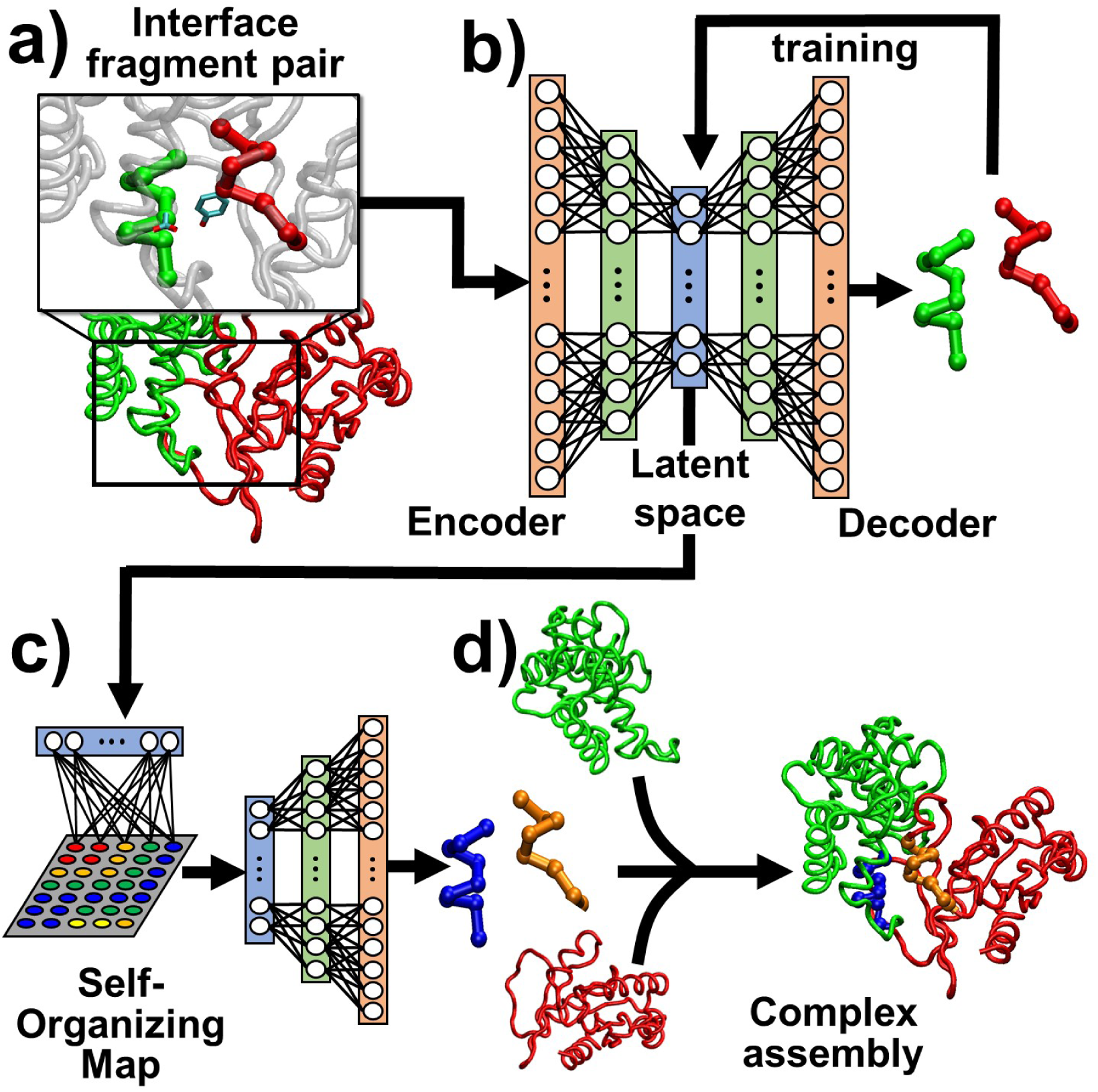
Given two interacting proteins, a pair of 9-amino-acid-long fragments is highlighted in **(a)** if the sidechains of their centered residues form at least one inter-molecular atomic contact. The structures of these interface fragment pairs were collected from a database of protein complexes and fed into an autoencoder. After training, the interface fragment pairs can be reconstructed from the output layer of the autoencoder, as shown in **(b)**. We further clustered the vectors in the latent space of the autoencoder by a self-organizing map. Highly clustered vectors were then sent to the latent space to generate new conformations of interface fragment pairs **(c)**. Finally, these newly generated interface fragment pairs were used to guide the assembly of protein complexes from separated monomers **(d)**.

Based on the above definition, we constructed a dataset of interface fragment pairs. We first selected protein complexes from the 3did database [30]. The database is categorized into groups of interacting domain pairs (IDPs). Each IDP further contains a large amount of 3D items which experimentally derived structures are available. Different 3D items within an IDP share the same Pfam index for both interacting domains. In order to reduce the sequence redundancy in our dataset, only one representative 3D item was selected from each IDP, which leads to a final selection of 4960 protein complexes. Given each protein complex, we first found out all the inter-molecular residue contacts. The list of all interface fragment pairs was then generated using the definition described in the last paragraph. By repeating this process for all 4960 protein complexes, a total number of 153127 interface fragment pairs were derived from the 3did database. Each item in the dataset contains two 9-amino-acid long fragments with their center residues forming an inter-molecular contact. Our dataset of interface fragment pairs kept the Cartesian coordinates of all these 153127 items.

### Design of autoencoder

An autoencoder neural network consists of an encoder and a decoder. The encoder learns to compress the input data from a higher dimension space to a lower dimension latent space, while the decoder tries to reconstruct the original data. It is an unsupervised learning algorithm that sets the target values to be equal to the inputs. Both the encoder and decoder in our model had a dense neural network architecture with one hidden layer. In order to reduce training complexity, the input, hidden, and output layers of the encoder and decoder were organized to mirror each other, as shown in **Fig 1b**. The number of neurons in both input and output layers equals 54, corresponding to the dimension of the conformational space for an interface fragment pair. The number of neurons in the hidden layers was set to 27, while the dimensions of the latent spaces were chosen to be equal to 18. All layers had a sigmoidal activation function and the mean square error was used as the loss function. The neural networks were trained by the backpropagation algorithm. Finally, training was carried out by using a batch size of *N* for 1000 epochs, in which *N* equals the number of interface fragment pairs in the training dataset. Within each epoch, samples were picked from the training set by random order.

During training, the coordinates of interface fragment pairs were fed to the input layer of the autoencoder. These coordinates were reconstructed from the output layer. By comparing the reconstructed coordinates to the native correspondence, the weights of neurons in all layers were then updated by the backpropagation algorithm. This process was iterated until the loss function was minimized. After training, the autoencoder was tested by reconstructing the structure of interface fragment pairs that were not in the training set. Moreover, vectors in the trained latent space were saved. They were projected onto a two-dimensional array using a self-organizing map algorithm. As a result, the decoder can be used to generate new conformations of interface fragment pairs by sampling the vectors in this new clustered space. Detailed process of SOM-based clustering is described below.

### Clustering of latent space by self-organizing map

In our SOM model, a set of neurons were arranged in a 100×100 rectangular array (**Fig 1c**). Each neuron has a vector of weights which length equals 18. This is equivalent to the length of vectors in the latent space. At the beginning of learning, the weights in all vectors were initialized to small random numbers. Within each learning step *t*, one vector in the trained latent space *x(t)* was randomly selected. The distances between the selected vector and the weight vectors of all neurons were then calculated. The neuron with the smallest distance was selected as winner. The weights of the winner *v* and its nearby neurons *k* were further updated towards the input vector *x(t)* by *Δw_k_(t)*. The weight update of neurons *k* can be expressed as *Δw_k_(t)=α(t)η(ν,k,t)[x(t)−w_ν_(t)]*. In this equation, *α(t)* is the learning rate which has a value between 0 and 1. It decreases monotonically along with the learning step *t*. Furthermore, *η(ν,k,t)* is a neighborhood function which value depends on the distance between the winner *v* and the selected neuron *k*. A Gaussian form was adopted to describe the neighborhood function. It can be written as *η(ν,k,t)=exp[−||r_ν_−r_k_||^2^/2σ(t)^2^]*, in which *r_v_* and *r_k_* denote the positions of neurons *v* and *k* in the 2D array, with *σ(t)* representing the effective range of the neighborhood, and is decreasing monotonically with the learning step *t*.

Above process was repeated for all vectors in the trained latent space with a total number of 20 cycles. The times of each neuron in the SOM that was selected as winners were recorded. As a result, the SOM algorithm clustered these vectors onto a two-dimensional map with preserved topology. Vectors that are similar to each other in the latent space have a tendency to be localized together by the neighboring neurons. The weights of these clustered neurons and times of being selected as winners can thus be used to reflect the overall distribution of vectors in the latent space. Neurons that have the highest chance of being selected as winners represent the most abundant vectors in the latent space. Their weights can be used to generate conformations of interface fragment pairs. More detailed strategy is described in the **Results**.

### Structural reconstruction of protein complexes by interface fragment pairs

Given a pair of interacting proteins I and J, we can reconstruct the structure of their complex from an interface fragment pair generated by the autoencoder. The detailed method is described as follows, as shown by the schematic in **Fig 1d**. Starting from the structure of two monomeric proteins, we assume any residue on their surfaces can interact to each other. A surface residue is defined as its solvent accessible surface area is larger than 10Å^2^. The solvent accessible surface area was calculated by the Shrake-Rupley algorithm using a water molecule probe with size of 1.4Å [31]. In order to explore all the possible binding modes for a given protein complex, we first enumerated all potential combinations of fragment pairs in both interacting proteins. A 9-amino-acid-long window was assigned to each monomer. The window can slide from the N to the C terminus along the primary sequence of a protein, while the nine consecutive residues in the window correspond to a fragment in its tertiary structure. As a result, the total number of fragment pair combinations for a given pair of interacting proteins with length *N_I_* and *N_J_* is (*N_I_* − 8)× (*N_J_* − 8).

For each combination, if the center residues in both fragments are on protein surfaces, we compared their structures to an autoencoder-generated interface pair consisting of fragments A and B. In order to do so, we simultaneously superimposed the residues in the window of protein I to the fragment A, and superimposed residues in the window of protein J to the fragment B. Similarly, we also superimposed residues in the window of protein I to fragment B, and superimposed window of protein J to fragment A. We calculated the root-mean-square-difference (RMSD) for both structural alignments. If the value of RMSD in either case was lower than 2Ǻ, we further superimposed the structure of the entire proteins I and J to their corresponding fragments A or B. After both monomers were superimposed to their corresponding interface fragments, a structural model of dimeric complex was constructed. The model was rejected if it contained inter-molecular clashes. The above process was iterated through all its (*N_I_* − 8)× (*N_J_* − 8) combinations for all interface fragment pairs generated from the autoencoder. This leads to an ensemble of reconstructed structural models of the protein complex. We have tested the likelihood of finding native-like structures of protein complexes in our reconstructed ensembles using a large-scale benchmark set.

## Results

### The autoencoder accurately reconstructed the conformation of interface fragment pairs

In order to train the autoencoder, we divided the 153127 interface fragment pairs in our dataset into a training set and a testing set. We randomly selected 1000 pairs from the dataset as the testing set, while the remaining ones were treated as the training set. Because the redundancy among protein complexes has been eliminated during the construction of our dataset, the sequence similarity of interface fragment pairs between the training and testing sets were thus minimized to the lowest level. The data format was further processed as follows. The structures of all interface fragment pairs in the dataset were aligned to the first one by the rigid body superposition. The cartesian coordinates of each fragment pair were then divided by the maximal coordinate value in the dataset. These scaled coordinates between 0 and 1 were used as input to the encoder.

During training, processed vectors of interface fragment pairs were fed to the input layer of autoencoder by random order. They were then reconstructed from the output layer. The difference between the reconstructed and the actual vectors were minimized through an iterative process as described in the method. The difference was calibrated as RMSD. We calculated the overall RMSD averaged by the entire training set after each run of iteration. The calculated RMSDs were plotted in **Fig 2a** as a function of the iteration. Three different designs of network topology were tested. The original network, denoted as ATE1, was introduced in the method. In addition, we also increase the number of neurons in the hidden layer from 27 to 36 in the second network, denoted as ATE2. Finally, we further add a second hidden layer in the third network, in which the first hidden layer contains 36 neurons and the second hidden layer contains 27 neurons. This is denoted as ATE3.

**Figure 2:**
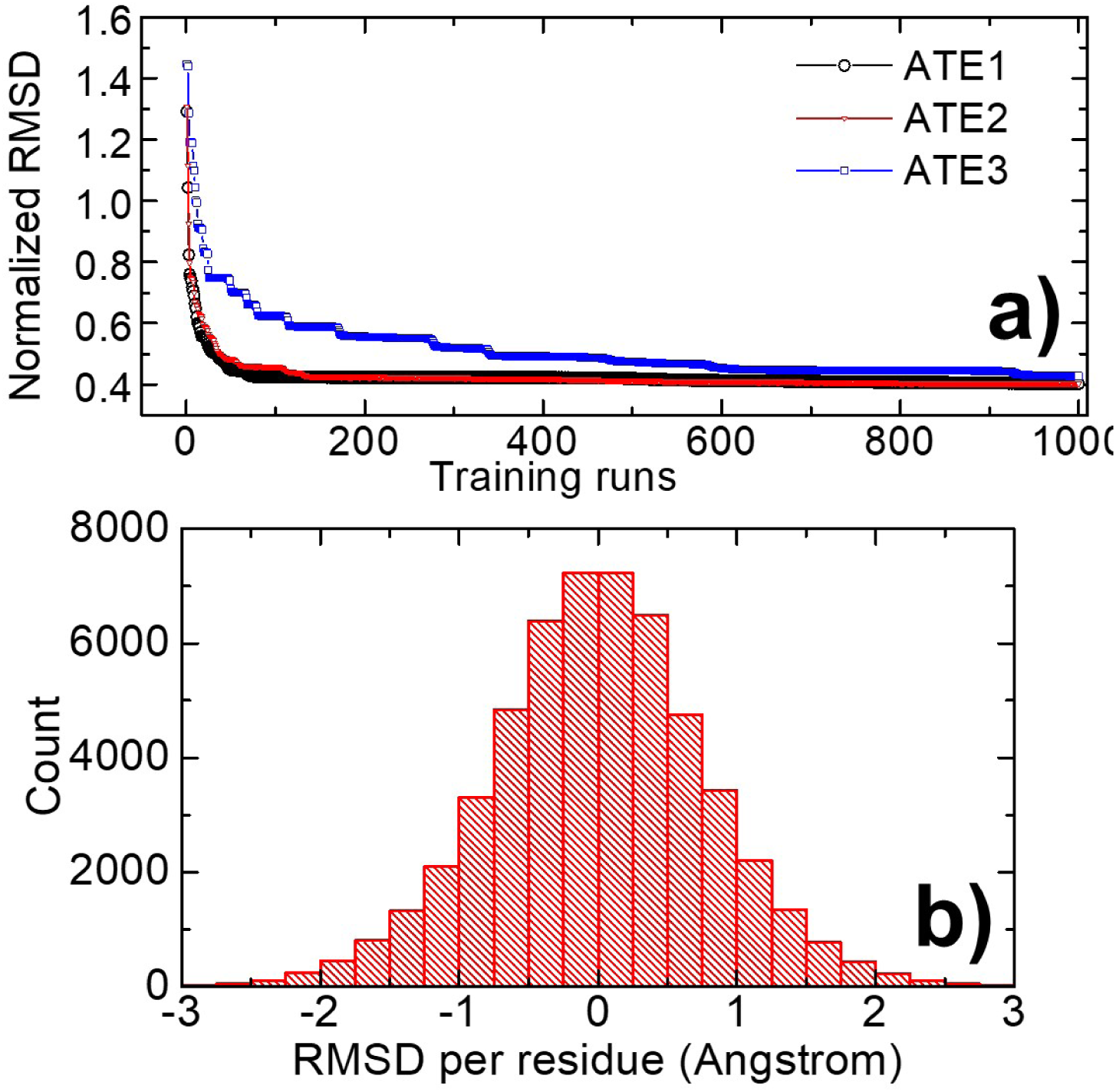
We designed three different network topologies in the autoencoder. The comparison of these three models during training is shown in **(a)**. The curves in the figure represent the overall RMSD averaged by the entire training set as a function of training runs. After training, the interface fragment pairs in the testing set were reconstructed. We compared the reconstructed results with their corresponding native conformations by calculating the RMSD of each residue in each aligned fragment. The RMSD distribution of all reconstructed interface fragment pairs was plotted as histograms in **(b)**.

The figure shows that in all three scenarios, the average RMSD of normalized vectors between the reconstructed and real systems dropped fast during the first 100 runs of training. After 100 runs, however, the decreases in RMSD became more slowly. This change in RMSD from a high level to a low level indicates the effectiveness of training. Moreover, we also compared the training results from different network topologies. Specifically, we found that changing the number of neurons in the hidden layer did not affect the training outputs. This is demonstrated in the figure by the fact that the results from ATE1, as shown by the black circles, are comparable to the results from ATE2, as shown by the red triangles. Adding an extra layer of hidden neurons in ATE3, however, made the training value of RMSD slightly higher than the first two models, as reflected by the blue squares in **Fig 2a**.

After training, the weights of all neurons in the autoencoder were fixed. The normalized vectors of 1000 interface fragment pairs in the testing set were fed to the input layer of encoder, while the vectors with the same length were then derived from the output layer of decoder. The output vectors were further scaled back into cartesian coordinates, so that the three-dimensional structures of interface fragment pairs were reconstructed. In order to test the accuracy of our reconstruction, we compared these reconstructed structures to their native conformations for all interface fragment pairs in the testing set. Given an interface fragment pair, we spatially aligned its reconstructed structure to the native conformation using rigid body superposition, and then calculated the RMSD of each residue in each aligned fragment. The calculations were carried out for all 1000 interface fragment pairs in the testing set. The distribution of all calculated RMSDs was plotted as histograms in **Fig 2b**. The standard deviation (SD) of the distribution is 0.8 Angstrom, while the largest RMSD is about 3 Angstrom. A majority of residues in reconstructed structures have RMSDs of less than 1 Angstrom from their corresponding native positions. This result indicates that the conformation of interface fragment pairs can be restored by the autoencoder with high fidelity.

**Fig 3** further shows the structural comparison of some interface fragment pairs selected from the testing set. The Cα atoms in each fragment are represented by beads, connected by a virtual backbone. Two fragments from the reconstruction are colored in orange and blue, while native pairs are colored in red and green, respectively. These selected fragment pairs represent diverse secondary structural elements observed at binding interfaces. In detail, **Fig 3a** gives an example of the packing between two interacting α-helices. In **Fig 3b**, the interface fragment pair contains two β-strands from the same piece of β-sheet that interact with each other through hydrogen bonds. **Fig 3c** shows a pair of structurally heterogeneous fragments: the interaction between an α-helix and a β-strand. Finally, in **Fig 3d**, two loops are co-localized at the binding interface. In all cases, the generated models are closely aligned to the native structures in three-dimensional space, suggesting that the autoencoder is able to rebuild the structure of any given interface fragment pair after training.

**Figure 3:**
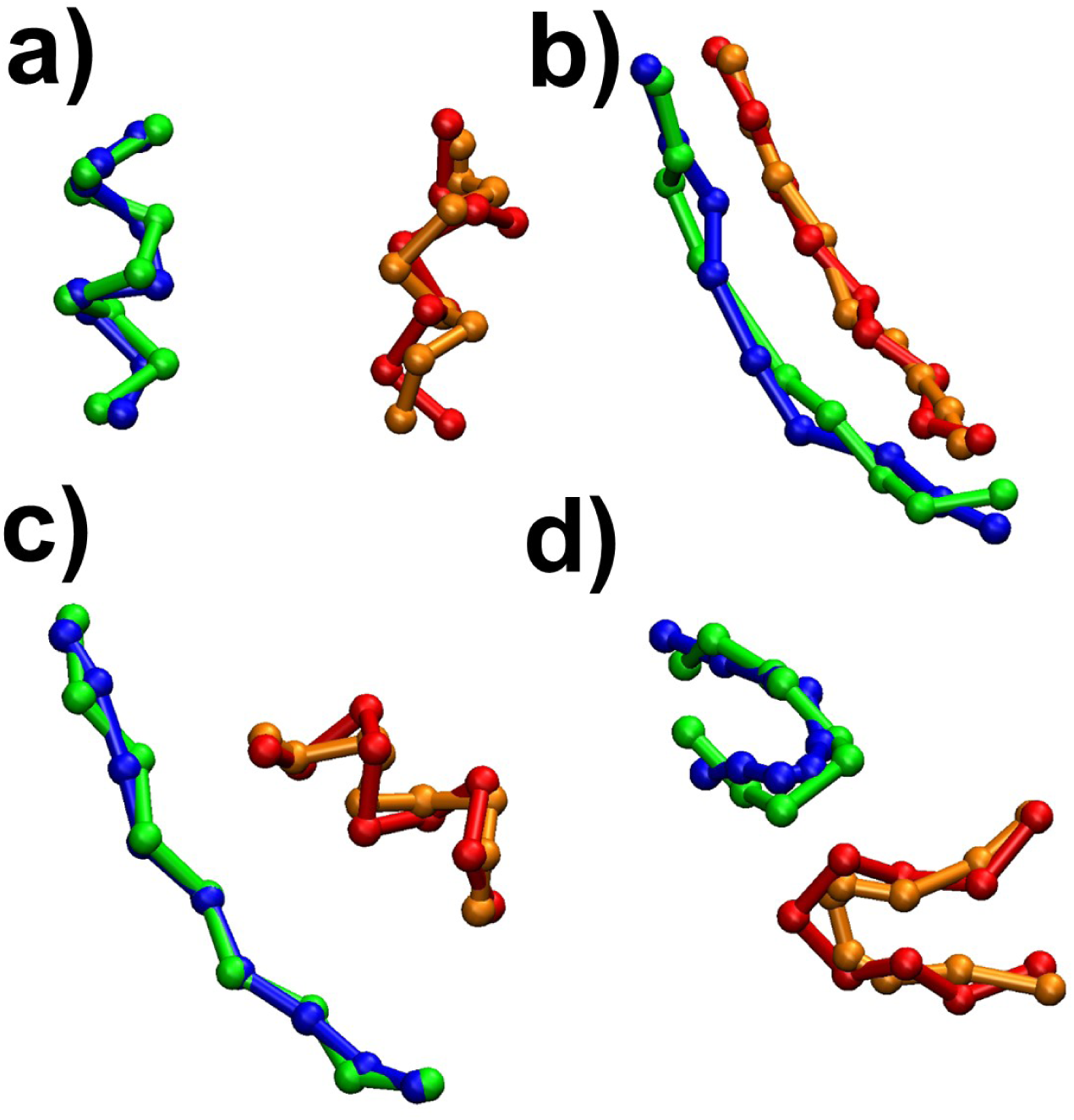
Some examples of our reconstructed interface fragment pairs are shown in the figure. They were aligned to their native structures for comparison. The reconstructed pairs are colored in orange and blue, while native pairs are colored in red and green, respectively. Specifically, **(a)** shows two fragments of α-helix packing against each other. In **(b)**, two fragments of β-strand interact with each other through hydrogen bonds. A pair of structurally heterogeneous fragments is shown in **(c)** The fragment pair contains an α-helix and a β-strand. Finally, in **(d)**, two loops are co-localized at the binding interface. In all cases, the generated models are closely aligned to the native structures.

### Vectors in the latent space of the autoencoder were clustered by a self-organizing map

The dimension of vectors in the latent space has already been reduced from the original degrees of freedom of an interface fragment pair. In order to characterize the correlation among these vectors, we further projected them onto a two-dimensional space by SOM. As described in the method, this two-dimensional space is discretized into a 100×100 array of neurons. The weight of each neuron is represented by a vector which length equals to the length of vectors in the latent space. The SOM model is based on an unsupervised learning strategy. In this strategy, all vectors in the latent space were thrown into the SOM space by random order. Each vector found a neuron in the space as winner based on their similarity. The weight of the winner neuron was then modified to resemble the selected vector. As a result, similar vectors in the latent space can be clustered into the neighboring neurons after multiple runs of feedback iterations. Moreover, the neurons that were most frequently selected as winners represent the most abundant vectors in the latent space.

Our SOM-based clustering results are visualized in **Fig 4a**. The array of neurons was plotted as a two-dimensional heat map. The map is from 1 to 100 along both dimensions, representing the number of neurons in the array. Different colors in the map indicate the number of each neuron that was selected as a winner during training. Red denotes the neurons that were most frequently selected as winners, while blue denotes the neurons that were least frequently selected as winners. The more quantitative index is coded by the color bar on the right side of the figure. In order to provide a closer-up view, some subsections of the map were further enlarged in the panels on the left side of the figure. The figure shows that, first, winners are not uniformly distributed. A majority of neurons have been selected as winners for a very few times, corresponding to the blue region in the map. On the other hand, there were 10% of neurons that were highly often to be selected as winners, representing over 50% of total vectors. Moreover, these small numbers of winner neurons were neither randomly scattered in the map nor aggregated together, but rather formed many micro-clusters. This result suggests that the vectors in the latent space not only have a high level of structural diversity, but also share some similarity within each small group.

**Figure 4:**
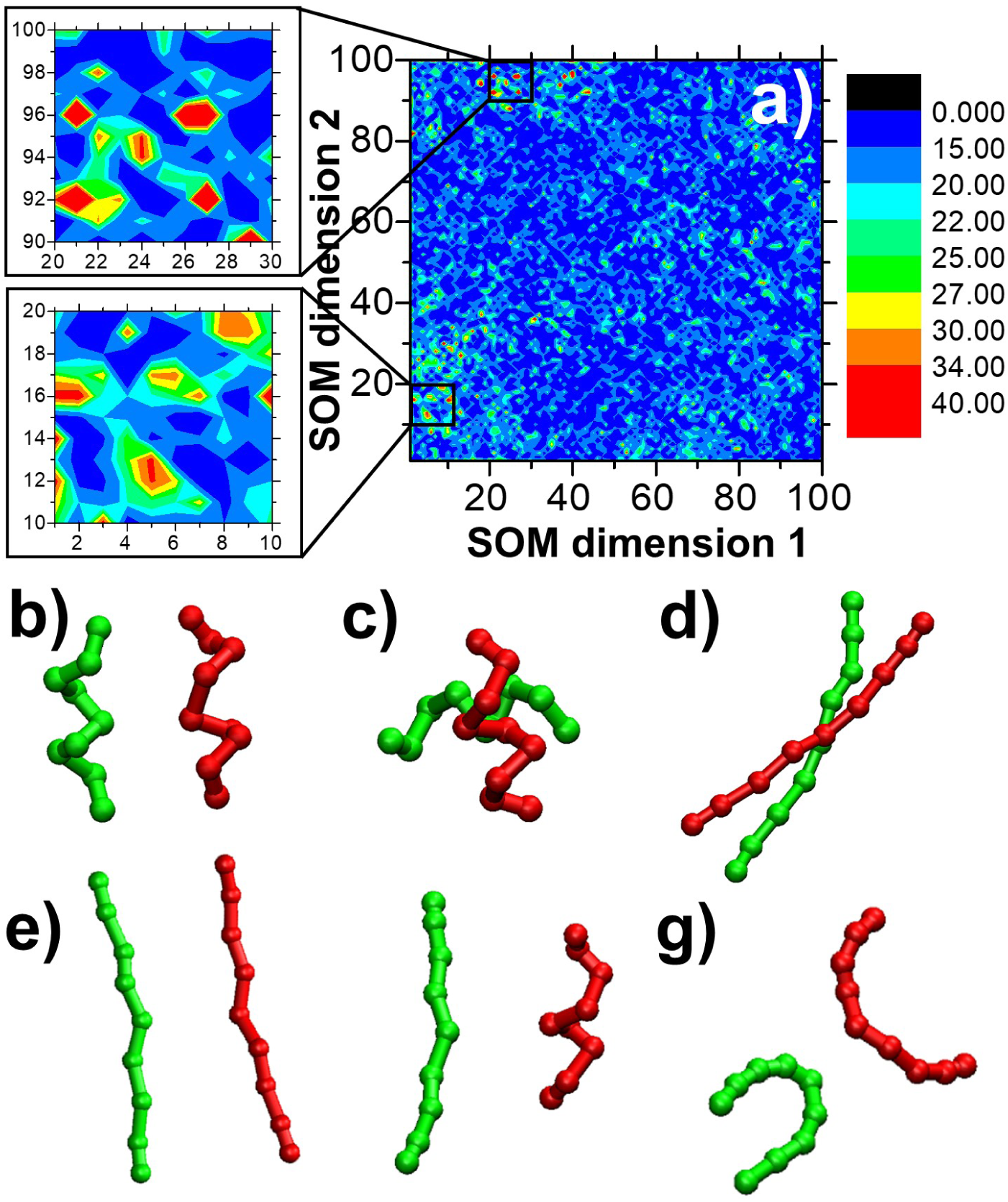
In order to characterize the correlation among vectors in the latent space, we clustered them together by self-organizing map. The clustered vectors were projected onto a two-dimensional space, as visualized in **(a)**. Different colors in the map indicate the number of each neuron that was selected as a winner during training. The color code is presented on the right side. Some subsections of the map are enlarged in the left panels to provide a closer-up view of the clustering result. After clustering, we generated new conformations of interface fragment pairs by sampling the neurons in the self-organizing map. We plotted come newly generated conformations, including the packing of two α-helices in **(b)** and **(c)**; the interaction between two β-strands in **(d)** and **(e)**; a heterogeneous pair formed between an α-helix and a β-strand **(f)**; and a pair consisting of two interacting loops **(g)**.

After the characterization of the vectors in the latent space, we further tested if new conformations of interface fragment pairs can be effectively generated by decoding the correlation of these vectors. Detailed strategy of conformational generation was guided by the SOM clustering results as follows. We sampled the two-dimensional space in our SOM model by randomly selecting the neurons in the map. The probability of a specific neuron being selected depends on how many times it became winners during training. The neurons that have never become winners had no possibility of being selected for generating new conformations. On the other hand, neurons that became winners more often had a higher possibility of being selected. After the random sampling, the weights of selected neurons were fed into the autoencoder as vectors in the latent space. Using previously trained parameters, the vectors were then passed through the decoder section. Finally, full-size coordinates in 54 dimensions were generated from the output layer. These coordinates were transferred back to the Cartesian system by multiplying the same value used in the previous encoding process.

We plotted the conformations of some interface fragment pairs that were newly generated through above sampling process. Some representative examples are shown from **Fig 4b** to **Fig 4g**. The Cα atoms in both fragments are represented by beads, connected by their corresponding virtual backbones. One fragment in these pairs is colored in red, and the other one is colored in green, respectively. We noticed that all the AI-generated interface fragment pairs in these figures are very similar to what can be found in native protein complexes. For example, both fragment pairs in **Fig 4b** and **Fig 4c** are formed between two α-helices. The packing of two interacting α-helices in these fragment pairs, however, is different. More specifically, the packing of two helical fragments in **Fig 4b** is parallel to each other, while the packing of two helical fragments in **Fig 4c** is orthogonal. These are the two most favorite modes of packing between α-helices in protein quaternary structural space. Two additional modes were observed in the generated fragment pairs in which β-strands were presented at the binding interface. They either interacted with each other in the same piece of β-sheet through hydrogen bonds, as shown in **Fig 4d**, or appeared in two separated β-sheets that faced each other, as shown in **Fig 4e**. Moreover, **Fig 4f** gives an example of a heterogeneous interface fragment pair that was generated between an α-helix and a β-strand, which has also been observed in the native protein complexes as shown in **Fig 3c**. Finally, two loops were observed at the binding interface in **Fig 4g**, similar to the native fragment pair shown in **Fig 3d**.

### Native-like protein complexes were assembled by the generated interface fragment pairs

In the last experiment, we tested how likely the interface fragment pairs generated from our autoencoder can be used to guide the assembly of protein complexes. Moreover, we want to inspect whether native-like conformations can be obtained by assembling a small subset of fragment pairs that most frequently appear at native protein interfaces. Therefore, only the neurons that formed large clusters in the SOM were selected. The weights of these neurons were fed to the latent space of our autoencoder. Using previously trained parameters, the vectors were passed through the decoder section, so that full-size coordinates in 54 dimensions were generated from the output layer. These coordinates were transferred back to the Cartesian system by multiplying the same value used in the previous encoding process. As a result, a total number of 777 interface fragment pairs were generated through this procedure.

In order to test if native conformation of protein complexes can be derived through these generated interface fragment pairs, we applied them to a large-scale protein-protein docking benchmark set [32] constructed by ZLAB which contains a set of 84 non-redundant protein complexes. For each complex in the benchmark, we first separated it into individual monomers. These monomers were then assembled back together by superimposing the structure of their corresponding fragments to each of the 777 generated interface fragment pairs, using the algorithm described in the method. Because the purpose of this test is to reconstruct protein complex from interface fragment pairs, but not to develop a new docking algorithm, we used bound structures of protein monomers during the test instead of unbound structures that are normally used to evaluate the performance of docking results. Consequently, a large ensemble of structural models was formed for each protein complex in the benchmark. Among the derived ensembles, we found the models which have the lowest RMSDs from the native structures of the respective protein complexes. The distribution of these RMSDs for all 84 protein complexes is shown in **Fig5a** as a histogram. For all cases, we found structural models which RMSDs are less than 6.0Å from their native structures. The average value of the distribution is 2.81Å. This result indicates that the native conformations of protein complexes were reproduced from our AI-generated interface fragment pairs with a high successful rate. It also suggests that structural space of protein-protein interactions can be encoded by only a limited number of binding modes.

In addition to comparing the best models in the benchmark test, we also evaluated how difficult to find any native-like conformations from individual ensembles. **Fig 5b** and **Fig 5c** show the probability plots of two protein complexes in the dataset. Each blue circle in the figures represents a structural model in the ensembles. They were ranked by their RMSD values along the x-axis. The y-axis indicates the percentile of finding structural models below specific RMSD. For example, 1% of structural models in the ensemble are below the RMSD of 2.5Å and 10% of structural models in the ensemble are below the RMSD of 5Å for the protein complex with PDB ID 1DE4, as shown in **Fig 5b**, with additional 40% of structural models below the RMSD of 7.5Å. The ensemble of the protein complex in **Fig 5c** (PDB ID 1F51) has the similar probability distribution. These figures confirmed that a relatively high amounts of structural models in our derived ensemble of protein complexes have native or near-native conformations.

**Figure 5:**
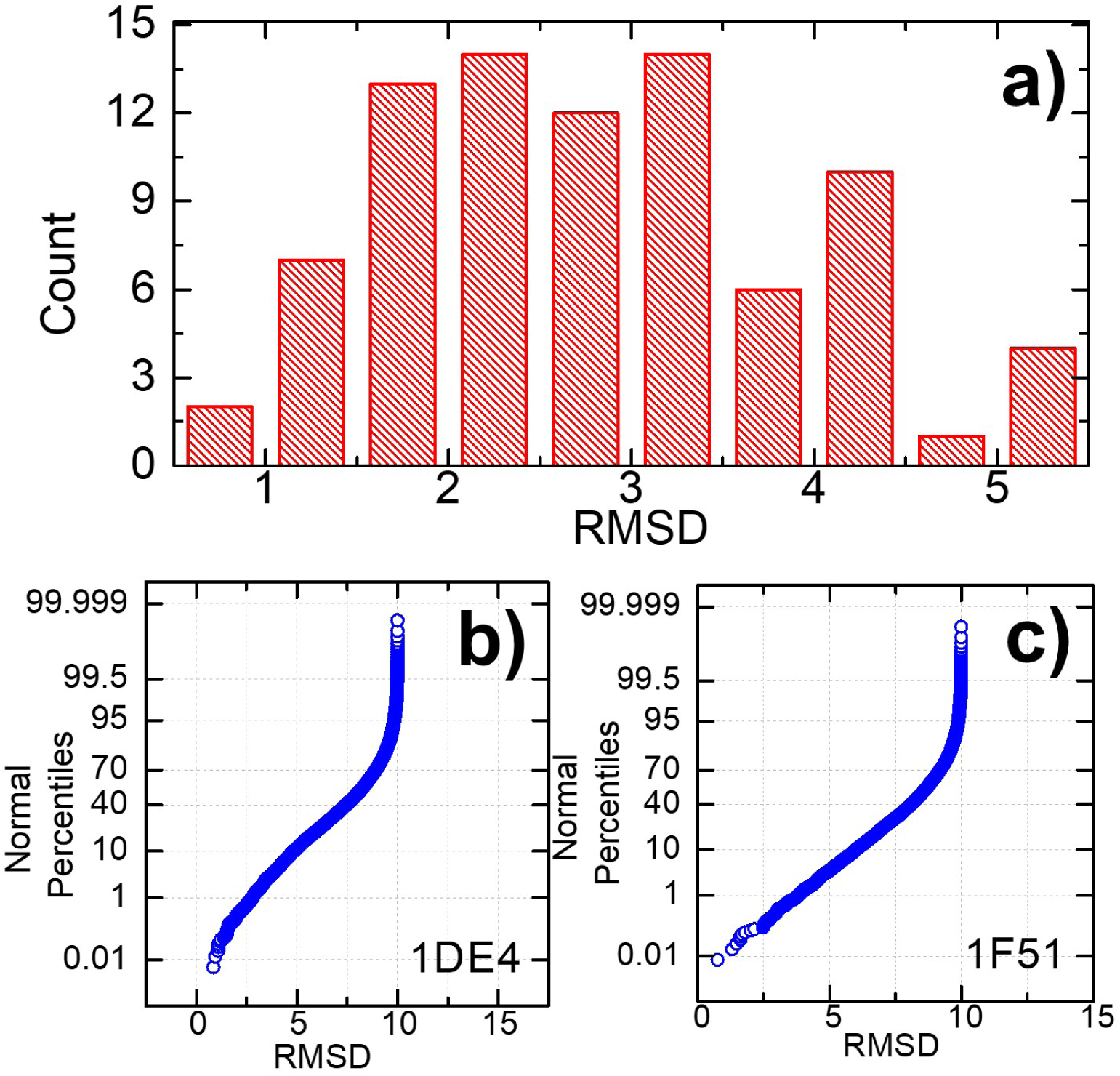
We tested if native conformation of protein complexes can be derived through these generated interface fragment pairs by a large-scale docking benchmark dataset. For each complex in the benchmark, a large number of structural models were generated. The models which have the lowest RMSDs from the native protein complexes were identified. The RMSD distribution of these best models is shown in **(a)** for all protein complexes in the benchmark. The probabilities of finding any native-like conformations have also been calculated for each protein complex in the dataset. Two specific examples are provided in **(b)** and **(c)**. Each blue circle in the figures represents a structural model in the ensembles. They were ranked by their RMSD values along the x-axis. The y-axis indicates the percentile of finding structural models below specific RMSD.

Some representative modeling results were selected in **Fig 6** from the benchmark. The Cα traces in orange and blue are the assembled structural models of two protein monomers which have the lowest RMSD. They were superimposed to their corresponding native structures that are colored in red and green with transparent cartoon representation. The interface fragment pairs used to assemble the complexes are highlighted by the beads in dashed yellow circles. The PDB ID and the RMSD value of each selected complexes are also listed below respective panel. **Fig 6a** shows a complex formed between a Fab fragment of a monoclonal murine IgG antibody and the major allergen from birch pollen Bet v 1 (PDB ID 1FSK). The fragment pair used to assemble the complex contains two β-strands at the binding interface. Similarly, **Fig 6b** shows a complex of the C-terminal half of gelsolin bound to monomeric actin (PDB ID 1H1V). The fragment pair used to assemble the complex contains two α-helices at the binding interface. We successfully found the native conformations of protein complexes in both cases. In contrast, **Fig 6c** and **Fig 6d** give two examples in which our best models are less similar to the native conformations. Specifically, the fragment pair used to assemble the complex in **Fig 6c** is formed between an α-helix from one protein and a β-strand from the other, while the fragment pair used to assemble the complex in **Fig 6d** contains two loops. In summary, these results suggest that homogenous fragment pairs between two α-helices or between two β-strands tend to have lower structural variations at binding interfaces than the heterogeneous fragment pairs or fragment pairs between two loops.

**Figure 6:**
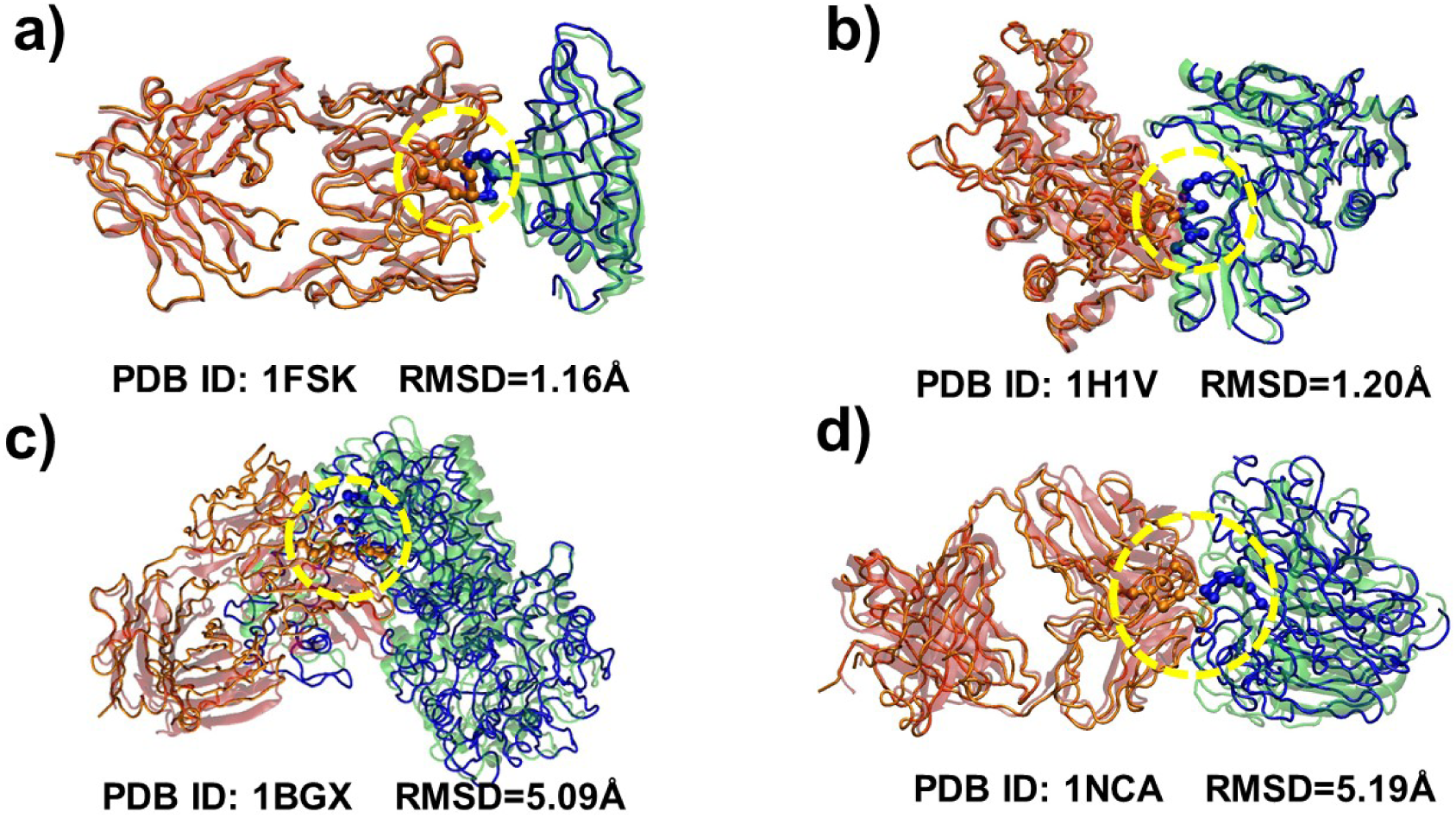
The structural models of our assembled protein complexes are compared to their native conformations. Two examples in which we successfully found the native conformations of protein complexes are shown in **(a)** and **(b)**. In contrast, **(c)** and **(d)** give two examples in which our best models are less similar to the native conformations. In all cases, the assembled structural models are represented by the Cα traces in orange and blue, while their corresponding native structures that are colored in red and green with transparent cartoon representation. The interface fragment pairs used to assemble the complexes are highlighted by the beads in dashed yellow circles. The PDB ID and the RMSD value of each selected complexes are listed below each respective panel.

## Concluding Discussions

The physical interactions between proteins dominate almost all biological processes in cells [33-35]. These interactions are largely determined by the structural properties of proteins at binding interfaces. However, only 6% of the known protein interactions have an associated experimental complex structure [36], which offers a large opportunity to computationally predict structural models of protein complexes. The recent development of deep learning computational methods has gained huge success when applied to modeling the structures of proteins and their interactions. The success of these applications indicates that common structural features exist at the binding interfaces of protein complexes, so they can be captured by artificial intelligence. In order to explore these features, we broke protein binding interfaces into pairs of interacting fragments. We collected a comprehensive set of interface fragment pairs, and encoded their structures into a low-dimensional latent space. Vectors in the latent space were clustered. New conformations of interface fragment pairs were then generated by decoding the vectors that are highly representative in the clusters. We showed that native-like protein complexes can be successfully assembled by only using a limited number of these newly generated interface fragment pairs. Our study, therefore, demonstrated that the structural space of protein-protein interactions is highly degenerate under the representation of interface fragment pairs. Features in this degenerate space can be well characterized by artificial intelligence.

Although the native-like conformations are among all the assembled structural models in most cases of the benchmark dataset, they cannot be identified from other models. This is due to the fact that the current model dose not incorporate sequence features of interface fragment pairs. Our primary interest is to analyze the structural features of protein complexes, but not to predict their structures. Nevertheless, the new method developed in this study will become practically useful after integrating the sequence features into our machine learning model. For instance, we can apply protein language models [37] to extract sequence features for all fragment pairs in the database. Moreover, currently available docking scoring functions [38, 39] can be applied to distinguish native-like protein complexes among all generated structural models. Additionally, protein binding site prediction [40, 41] can also be integrated into our method so that only the sequences close to these predicted binding sites in two interacting proteins will be selected to generate the structure models of interface fragment pairs. This could further narrow down the search space when we generate structural ensembles of target complexes. In summary, the interface fragment pairs generated by artificial intelligence offer an efficient tool for modeling the structure of protein complexes.

## ACKNOWLEDGMENTS

This work was supported by the National Institutes of Health under Grant Numbers R01GM120238 and R01GM122804. The work is also partially supported by a start-up grant from Albert Einstein College of Medicine. Computational support was provided by Albert Einstein College of Medicine High Performance Computing Center.

## AUTHOR CONTRIBUTIONS

Z.S and Y.W. designed research; Z.S. and K.D. performed research; Z.S., K.D., and Y.W. analyzed data; Y.W. wrote the paper.

